# Airway macrophage specific expression of a dense glycocalyx and its remodelling following viral infection

**DOI:** 10.1101/2025.11.20.689471

**Authors:** Ziyun Zhang, Iashia Z Nabi, Tracy Hussell, Douglas P Dyer

## Abstract

The glycocalyx is best studied for its regulation of immune cell trafficking from the vasculature into inflamed tissues. The glycocalyx is composed of a range of glycans and glyco-proteins that form a peri-cellular matrix and its shedding into the blood is observed in a variety of inflammatory conditions, including viral infections. We now report that macrophages express a glycocalyx that is dependent on their anatomical location and the mediators driving their differentiation from monocytes. Furthermore, during inflammatory conditions caused by viral infection, the macrophage glycocalyx is remodelled in complex ways. Overall, our study provides a novel pathway involved in macrophage inflammatory responses, through remodelling of the glycocalyx halo expressed at steady state, and how this is restored in repair, particularly at mucosal tissue sites.

## Introduction

The glycocalyx is a meshwork of different glyco-conjugates that forms a pericellular matrix on the surface of most cells^1^. Its function ranges from barrier formation to transduction of signals from the extracellular environment^2^. This meshwork is most studied on endothelial cells where it regulates cell extravasation through steric hindrance of homing receptors and/or by electrostatic repulsion and vascular homeostasis^3^. Proteoglycans are key components of the glycocalyx and are composed of protein cores with sulfated glycosaminoglycan (GAG) sugar side chains (e.g. heparan sulfate (HS) and chondroitin sulfate (CS)) and the unsulfated GAG hyaluronan (HA)^4^. Proteoglycans that tether the glycocalyx to the cell surface include syndecans (SDC) 1-4. These transmembrane tethers provide a means for the cell to sense their local microenvironment via the glycocalyx in addition to their role as receptors for chemokines^5–8^.

More recently, a glycocalyx has been described on other cell types, including macrophages where it regulates efferocytosis of apoptotic cells as well as on monocytes where it regulates their migration into tissues^9–11^. The glycocalyx on airway epithelial cells is shed on injury^12^, and in response to LPS^13^, influenza virus^14^ and bleomycin^15^, leading to barrier dysfunction, epithelial apoptosis, surfactant dysfunction, impaired epithelial repair and susceptibility to secondary bacterial pneumonia^14^. The method of glycocalyx shedding seems to depend on location. Small fragments, like HS, are released into the blood by heparanase, whereas larger fragments in the airway indicate shedding of the tethering proteoglycans, such as SDC-1^16,17^.

The heterogeneity of glycocalyx on cells in different anatomical locations is largely untested, but has recently been explored in the endothelial compartment across tissues^18,19^. Here, we test the hypothesis that the lung macrophage glycocalyx is context-specific. Limitation of macrophage reactivity in the airspaces is important to prevent overt inflammation to innocuous antigens or commensal microbes^20^. Numerous site-specific mechanisms exist that raise the threshold for airway macrophage activation. These include ligation of airway macrophage CD200 receptor by epithelial expressed CD200^21^, lung epithelial-derived surfactant proteins binding and blocking macrophage toll-like receptors^22^ and enhanced concentrations of airway IL-10, GM-CSF and activated TGFβ, and many more^23^.

We now show that a distinct glycocalyx layer surrounds airway, but not lung interstitial, macrophages at steady state, which we reproduce *in vitro* by differentiating monocytes into macrophages with GM-CSF (but not M-CSF), followed by further polarisation with IL-4. Interestingly, interstitial macrophages migrating into the lung during inflammation gain a dense glycocalyx, as do transferred monocyte-derived macrophages. This demonstrates that the airway microenvironment facilitates the acquisition of a glycocalyx on macrophages, which potentially regulates their function.

## Results

The relative abundance of glycocalyx on lung macrophages occupying different anatomical compartments is unknown. Fluorescently labelled Wheat Germ Agglutinin (WGA) that binds glycoconjugates (N-acetylglucosamine and sialic acid residues) on cell membranes, was used to visualise the macrophage glycocalyx in lung paraffin-embedded tissue sections^24,25^. A dense glycocalyx was observed on macrophages in the airway lumen adjacent to the airway epithelium (Figure 1 A-D). By contrast, CD64+ macrophages in the lung interstitial tissue showed little binding to WGA (Figure 1 A, E-G). A similar result was also obtained after retrieving airway macrophages and interstitial macrophages by tissue digestion and analysing glycocalyx expression by flow cytometry (Figure 1 H, I) distinguished based on the staining protocol shown in supplementary figure 1. Macrophages expressing WGA-binding glycans in the airways (but not the interstitial macrophages) were confirmed in a separate experiment as double positive for CD64 and Siglec F (Figure 1 J).

**Figure 1:**
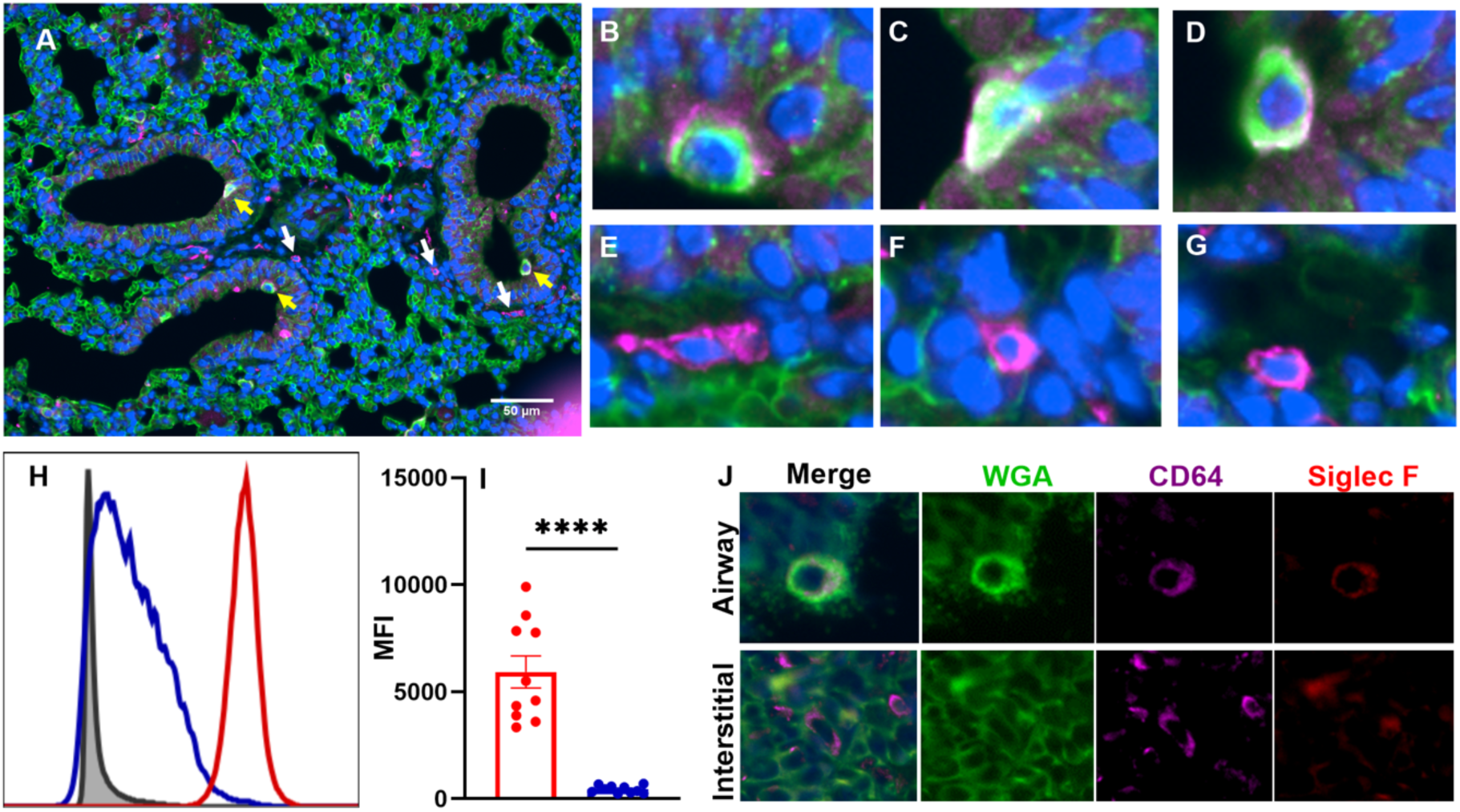
Macrophages in the lung and airway express the glycocalyx. Representative Confocal images of a healthy mouse’s lung showing general glycocalyx (WGA lectin: green), macrophages (CD64: Cyan), and DNA material (DAPI: Blue). Scale bar: 50μm. Macrophages are located in the airway (yellow arrows) and in the tissue (white arrows) (**A**). Detailed views of airway macrophages (**B, C, and D**). Detailed views of tissue macrophages (**E, F, and G**). Cells were isolated from the digested lungs of female C57BL/6 mice to identify alveolar macrophages, and interstitial macrophages. Histogram comparison of fluorescence signals from WGA lectin from alveolar macrophages (red), interstitial macrophages (blue), and fluorescence minus one control (FMO) (grey) (**H**). The medium fluorescence intensity (MFI) for WGA lectin of alveolar macrophages (red) and interstitial macrophages (blue) (**I**). Representative confocal images of healthy naïve mouse lungs showing general glycocalyx (WGA lectin: green), macrophage marker (CD64: cyan), and alveolar macrophage marker (Siglec-F: red) (**J**). Each dot represents an individual mice with data pooled from three independent experiments. Data are plotted as the mean ± SEM and analysed using unpaired t-tests. ****, P ≤ 0.0001.

To understand what factors might drive glycocalyx expression on airway macrophages (and whether they produce it themselves), we differentiated bone marrow mesenchymal stem cells into macrophages (BMDMs) using GM-CSF or M-CSF (Figure 2 A). Both GM-CSF and M-CSF-derived BMDMs bound highly to WGA, and expressed SDC-1 and CD44 by flow cytometry analysis (supplementary Figure 2). However, RT-qPCR revealed that RNA levels for some genes involved in glycocalyx synthesis and regulation were significantly greater in GM-CSF-derived, compared to M-CSF-derived BMDMs. This included raised mRNA for *has-1, sdc-2* and *mmp-9, −12, and −13* when presented as a fold change of GM-CSF/M-CSF BMDMs (Figure 2 B-O) or as relative expression levels over the housekeeping gene *b2m* (supplementary Figure 3).

**Figure 2:**
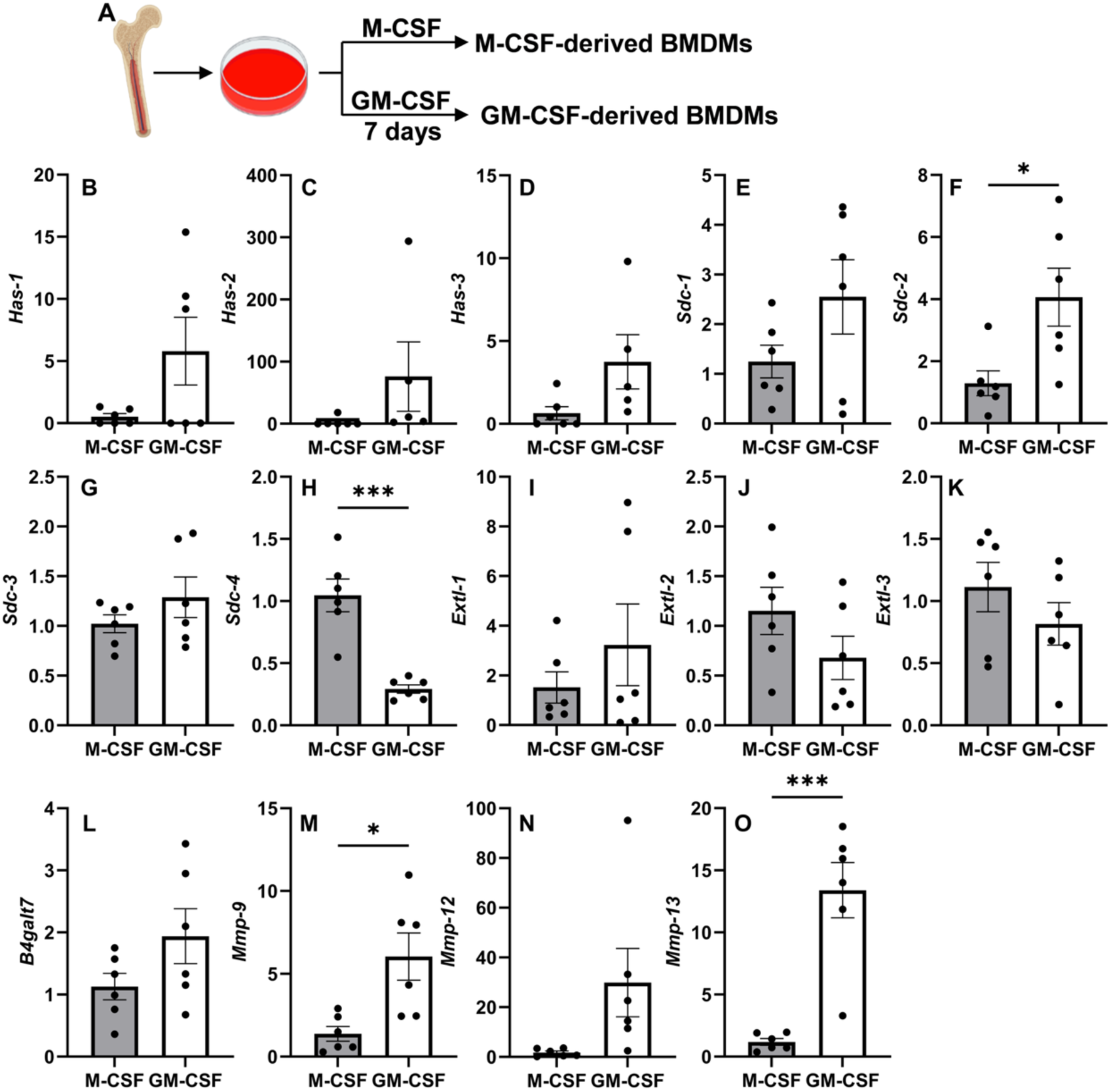
GM-CSF-derived BMDMs expressed higher levels of genes for glycocalyx degradation than M-CSF-derived BMDMs. Bone marrow cells were derived with M-CSF or GM-CSF (**A**). The glycocalyx-relative mRNA expressions were measured in GM-CSF- and M-CSF-differentiated BMDMs. RT-qPCR data are normalised to the housekeeping genes *b2m* and displayed as fold change over the M-CSF-derived BMDMs. *Has-1* (**B**), *has-2* (**C**), *has-3* (**D**), *sdc-1* (**E**), *sdc-2* (**F**), *sdc-3* (**G**), *sdc-4* (**H**), *extl-1* (**I**), *extl-2* (**J**), *extl-3* (**K**), *b4galt7* (**L**), *mmp-9* (**M**), *mmp-12* (**N**), and *mmp-13* (**O**). Each dot represents a technical repeat, data are representative of two independent experiments, plotted as the mean ± SEM. and analysed using unpaired t-tests. *, P ≤0.05; **, P ≤ 0.01; ***, P ≤ 0.001.

We next determined whether further polarisation of M-CSF-derived BMDMs with IFN-ƴ/LPS or IL-4 (Figure 3 A) affected glycocalyx expression. The effectiveness of polarisation was determined by the relative mRNA expression of *nos-2* and *chi3l-1*, which, as expected, were raised in M1 and M2 polarising stimuli^26^, respectively (Supplementary Figure 4). No difference in the already high levels of WGA binding was observed between the different conditions (Figure 3 B). In M-CSF-derived macrophages, SDC-1 was low in non-polarised macrophages and those polarised with IFN-ƴ/LPS, but significantly greater expression occurred in the presence of IL-4 (Figure 3 C). No difference was observed in the low levels of HS when analysed by flow cytometry (Figure 3 D). For GM-CSF-derived BMDMs, no difference was observed for WGA, SDC-1 or HS levels (Figure 3 E-G).

**Figure 3:**
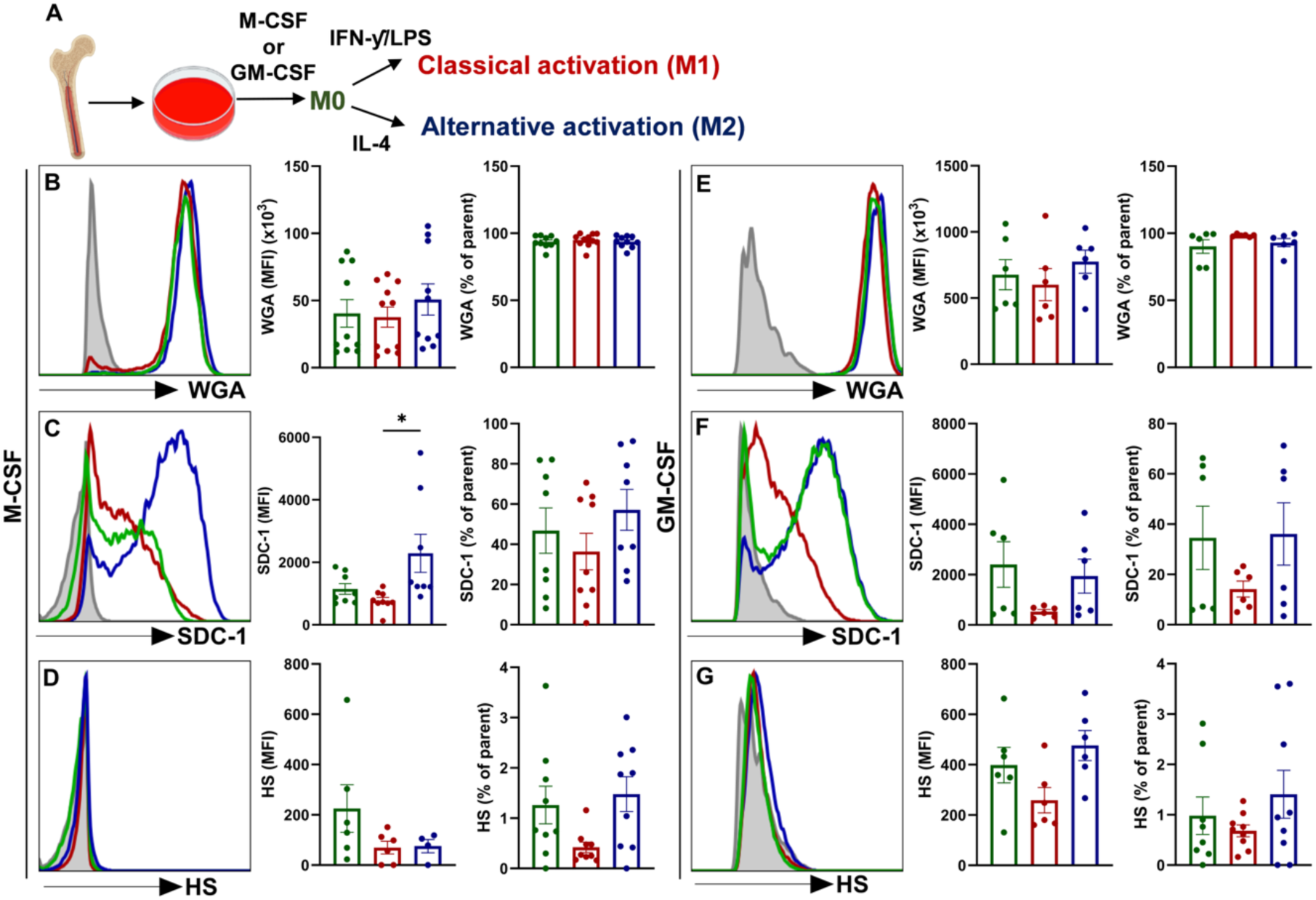
Classical or alternative activation of M-CSF and GM-CSF-derived BMDMs subtly regulates their glycocalyx content. The femurs of C57BL/6 female mice were flushed with HBSS to collect bone marrow cells. The bone marrow cells were then cultured with 20 ng/ml M-CSF, or GM-CSF for 7 days. After 7 days, the cells were left unstimulated (M0) (green) or polarised to classically (M1) (red) and alternatively activated (M2) (blue) macrophages (**A**). The representative histograms, MFI and percentage of M-CSF-derived BMDMs positive for glycocalyx as indicated by WGA lectin (**B**), SDC-1 (**C**), and HS (**D**). The MFI and percentage of GM-CSF-derived BMDMs positive for glycocalyx as indicated by WGA lectin (**E**), SDC-1 (**F**), and HS (**G**). Each dot represents a technical repeat, data are representative of two independent experiments, plotted as the mean ± SEM and analysed using an ordinary one-way ANOVA and Tukey’s multiple comparisons tests. *, P ≤ 0.05.

M-CSF-and GM-CSF-derived BMDMs, with IFN-ƴ/LPS or IL-4 polarisation, were lysed to examine the mRNA of glycocalyx modifiers, which was plotted as relative expression (fold changes) of the unpolarised BMDMs (Figure 4). Relative expression levels in comparison to the housekeeping gene *b2m* are shown in Supplementary Figure 5. The expression of glycocalyx modifiers altered with different stimuli. For example, IFN-ƴ/LPS induced *has-1* transcription (Figure 4 A) but reduced *has-2* transcription (Figure 4 B) and IL-4 induced *has-3* transcription (Figure 4 C). Furthermore, *sdc-1, −2 and −3* expression was reduced in response to IFN-ƴ/LPS (Figure 4 D-F), while the expression of *sdc-4* was enhanced by IL-4 (Figure 4 G). IFN-ƴ/LPS also caused reductions of *extls* and *b4galt7* that drive the biosynthesis of HS (Figure 4 H-K)^27^. Therefore, stimuli driving type 1 cytokines inhibited the majority of assessed glycocalyx synthesis-related gene expression in macrophages. *Mmp-9*, *-12*, and *-13* were induced by the IL-4 polarisation (Figure 4 L-N). Despite the fold changes, the relative amount of transcription of *has*, *sdc* and *extl* families of genes were low in comparison to the amount of the housekeeping gene *b2m* (Supplementary Figure 5).

**Figure 4:**
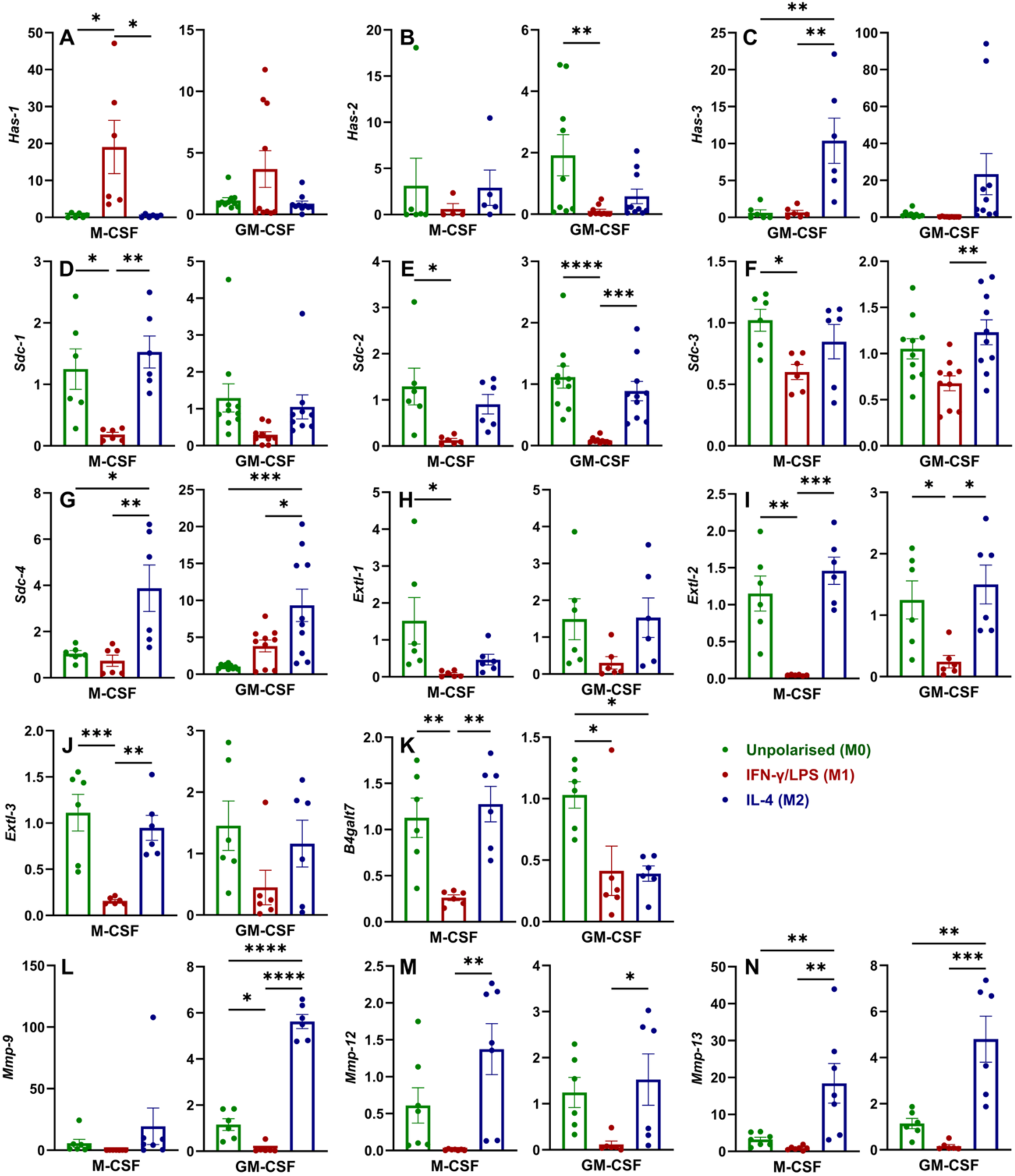
Classical activation of BMDMs reduced the transcription of most glycocalyx-related genes. M-CSF-derived BMDMs and GM-CSF-derived BMDMs were left unstimulated (green) or polarised to classically (M1) (red) and alternatively activated (M2) (blue) macrophages. The relative mRNA expressions were measured and displayed as fold change over the level of unpolarised BMDMs (set as 1). *Has-1* (**A**), *has-2* (**B**), *has-3* (**C**), *sdc-1* (**D**), *sdc-2* (**E**), *sdc-3* (**F**), *sdc-4* (**G**), *extl-1* (**H**), *extl-2* (**I**), *extl-3* (**J**), *b4galt7* (**K**), *mmp-9* (**L**), *mmp-12* (**M**), and *mmp-13* (**N**). Each dot is a technical repeat, data are representative of two independent experiments, plotted as the mean ± SEM and were analysed using unpaired t-tests. *, P ≤ 0.05; **, P ≤ 0.01; ***, P ≤ 0.001; ****, P ≤ 0.0001.

The reduction of the macrophage glycocalyx by type ! cytokines stimuli was confirmed using Poly I:C (a viral mimetic that induces a type-1 inflammatory response^28^) as the stimulus for the alveolar macrophage MH-S cell line, where WGA glycocalyx, SDC-1 and HA (Figure 5 A-C) was reduced on the cell surface. In the cell culture supernatant, we also observed a greater presence of soluble HS that has been released from Poly IC treated MHS cells (Figure 5 D), but not HA or SDC-1 (Figure 5 E, F).

**Figure 5:**
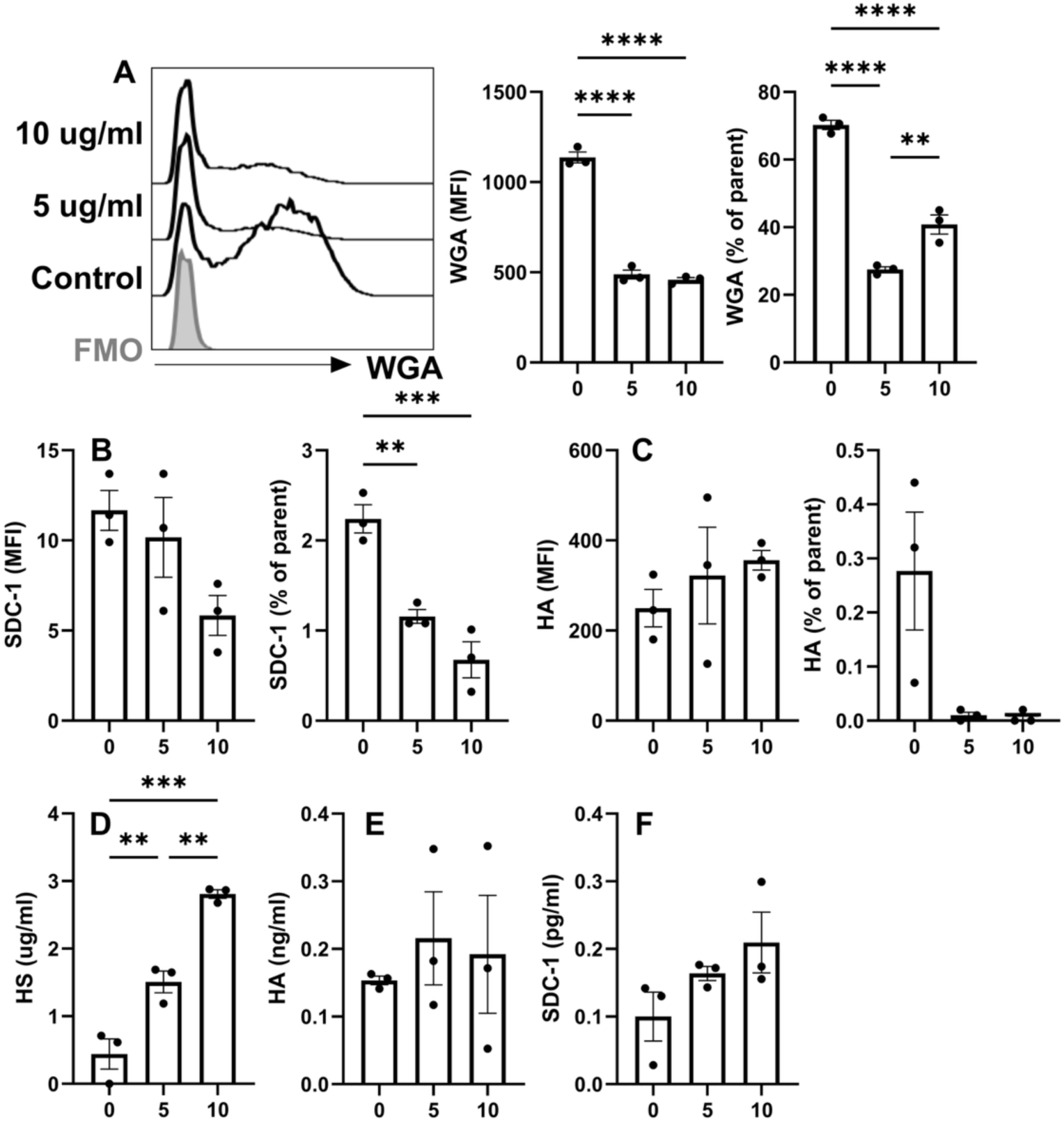
Incubation of MH-S cells with Poly I:C reduces the glycocalyx on the cell surface. MH-S cells were incubated with 5 ug/ml or 10 ug/ml Poly I:C for 24 hours or tissue culture medium alone. Representative histogram of the lectin WGA staining: Poly I:C treatment of 10 ug/ml, 5 ug/ml), medium-only control, and FMO control (grey). MFI and percentage of positive cells of WGA lectin staining (**A**). Components of glycocalyx are shown by SDC-1 (**B**), and HA (**C**). The cell culture supernatant was collected after 24 hours of stimulation and tested by ELISA for HS (**D**), HA (**E**), and SDC-1 (**F**). Each dot represents a technical repeat, data are plotted as the mean ± SEM and were analysed using an ordinary one-way ANOVA and Tukey’s multiple comparisons tests. *, P ≤ 0.05; **, P ≤ 0.01; ***, P ≤ 0.001; ****, P ≤ 0.0001.

Since our *ex vivo* and *in vitro* studies suggest glycocalyx is reduced in the presence of type 1 cytokines, we next examined macrophages in an *in vivo* murine model of lung influenza virus infection. At day 7 after infection, the weight loss is pronounced (Figure 6 A). Myeloid cells were gated as shown in Supplementary Figure 6 A. Airway macrophages maintained a high glycocalyx level, as shown by their WGA binding level irrespective of influenza virus infection (Figure 6 B), whereas they up-regulated SDC-1 (Figure 6 C). Interestingly, interstitial macrophages up-regulated both the WGA staining (Figure 6 D), and SDC-1 after influenza virus infection (Figure 6 E). We also observed some differences in WGA, but not SDC-1, on other cell subtypes (Supplementary figure 6 B-G). We next allowed the mice to recover from influenza virus infection and harvested tissues at day 14 post-infection (Figure 6 F). Again, at this time point, we observed an up-regulation of glycocalyx (WGA binding) and SDC-1 on interstitial macrophages (Figure 6 G, H). This suggests that, despite type 1 inflammatory cytokines reducing macrophage glycocalyx *in vitro*, the airway microenvironment *in vivo* overrides this and drives high macrophage glycocalyx levels even in the presence of viral triggers.

**Figure 6:**
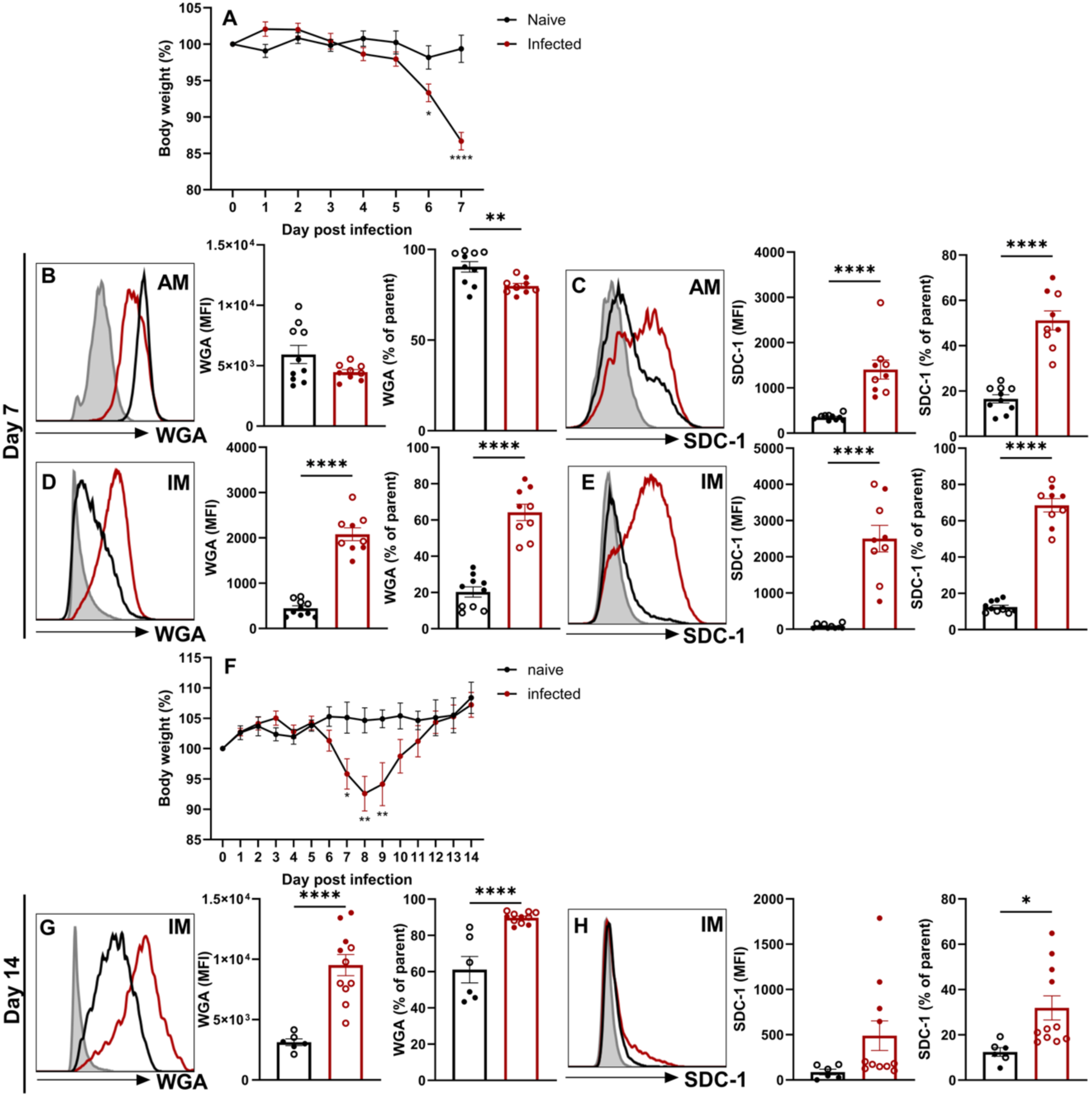
Influenza virus infection promotes SDC-1 expression on both alveolar macrophages and interstitial macrophages. At day 0, mice were intranasally given 5 PFU IAV PR8, or given PBS in the naïve control group. On day 7, the lung tissue from both groups was harvested. The tissue was digested and stained for flow cytometry analysis. Average weight change of naïve and infected mice from day 0 to day 7 (**A**). The MFI, and percentage, of general glycocalyx were shown by WGA lectin staining of AMs (**B**) and IMs (**D**) on day 7. The medium fluorescence intensity (MFI), and percentage, of SDC-1 of AMs (**C**) and IMs (**E**) on day 7. The influenza virus infection experiment was repeated, and the mice were harvested on day 14. Average weight change of naïve and infected mice from day 0 to day 14 (**F**). The MFI and percentage of WGA lectin (**G**), and SDC-1 (**H**) of IMs on day 14. Naive (orange), infected (red), and FMO control (blue). Each dot represent an individual mouse, data are pooled from two independent experiments, are presented as mean ± SEM and were analysed using unpaired t-tests with Welch’s correction. *, P ≤ 0.05; **, P ≤ 0.01; ***, P ≤ 0.001; ****, P ≤ 0.0001. Data from the two experiments were distinguished by the shape of the points.

We next directly tested whether the airway microenvironment drives high levels of macrophage glycocalyx. To do so, we collected bone marrow from CD45.1^+^ PeP-3 mice, differentiated them *in vitro* into macrophages with M-CSF (which are glycocalyx low) and transferred 1×10^6^ cells intranasally into CD45.2^+^ C57Bl/6 mice (Figure 7 A). Bronchoalveolar lavage (BAL) fluid and lung tissue were harvested 7 and 14 days later. The CD45.1^+^ transferred macrophages were clearly visible amongst the host CD45.2^+^ macrophages in the lung and airway. Local AMs and IMs were distinguished based on the expression of Siglec F. BAL does not remove the contents of the entire airways as the fluid doesn’t reach all of the alveolar sacs. Therefore, the digested lung always contains a proportion of residual airway macrophages, which are easily distinguished by flow cytometry. As expected, few Siglec F^-^macrophages were isolated from the BAL, thus the transferred cells collected from the BAL fluid were only compared with Siglec F^+^ airway macrophages (Figure 7 B). We compared WGA intensity among transferred CD45.1^+^, original local airway, and interstitial, macrophages from digested lung tissue. In the lung tissue at day 7, the greatest expression was observed on original local airway macrophages that had not been removed by BAL (Figure 7 C). However, by day 14, both local and transferred macrophages in the lung expressed equivalent levels of WGA (Figure 7 D). This was also observed in cells recovered by BAL, where at day 7 there was still a lower level of glycocalyx on CD45.1^+^ transferred macrophages, compared to local AMs (Figure 7 E). Again, by day 14 transfered macrophages had accumulated comparable glycocalyx levels (WGA^+^) to the local AMs (Figure 7 F). Furthermore, as expected, transferred macrophages reaching the airway by day 7 of infection had lower Siglec F expression compared to original airway macrophages. This also increased in proportion by day 14 (Figure 7 G, H), demonstrating development of the transferred cells into an AM-like phenotype. By focussing on the population of transferred macrophages that expressed higher levels of Siglec F we were able to demonstrate that these bound greater levels of WGA (indicative of glycocalyx) (Figure 7 I, J). Therefore, transferred cells enter the lung and airways and change their expression of glycocalyx, possibly via regulation of tethering receptors such as syndecan 1^29^.

**Figure 7:**
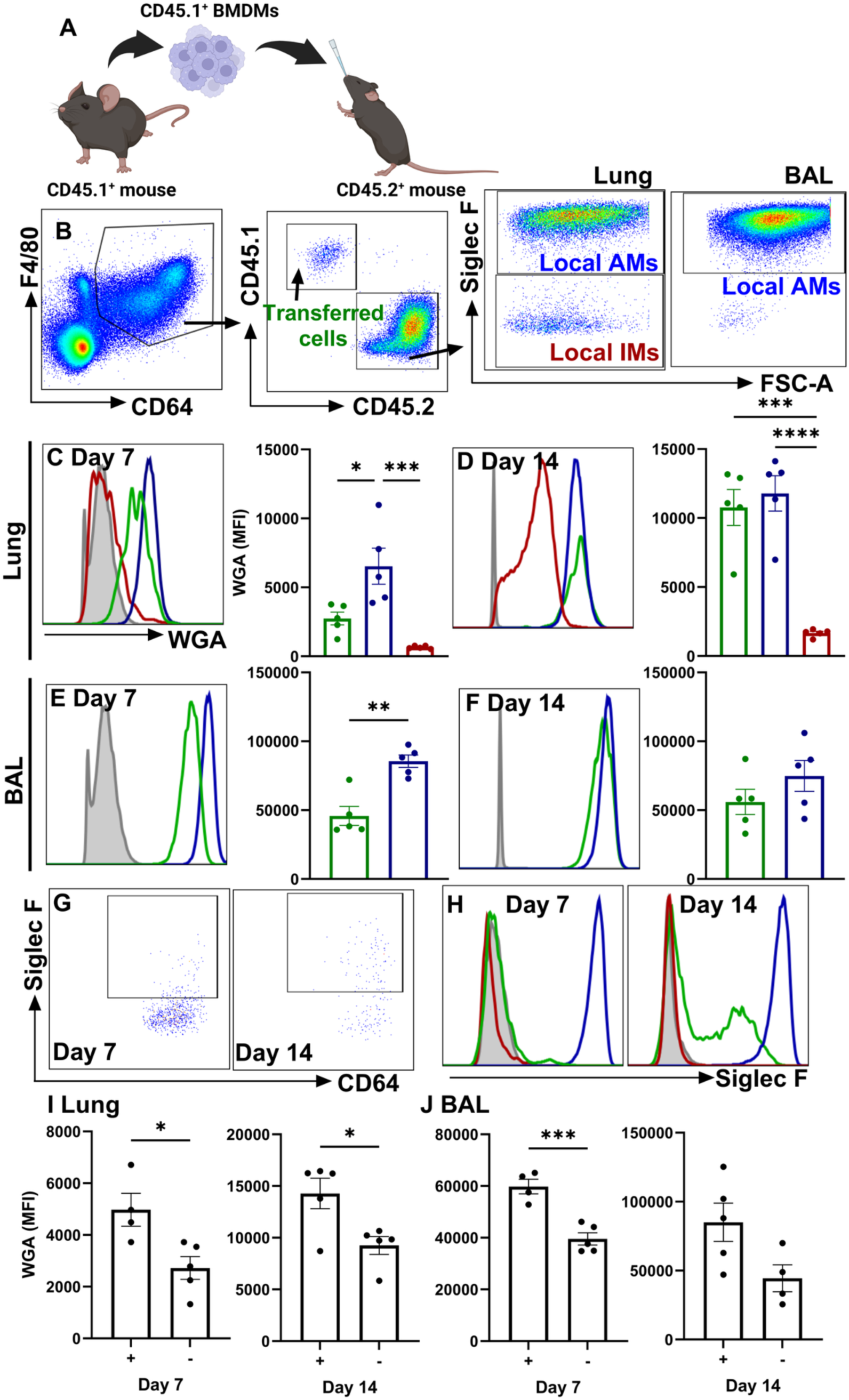
BMDMs transferred to the lung increased glycocalyx levels from day 7 to day 14. BMDMs from CD45.1^+^ mice were differentiated into macrophages with M-CSF and intranasally instilled in CD45.2^+^ wild-type C57BL/6 mice (**A**). At 7- or 14-day post-instillation, the BAL fluid and lungs from recipient mice were collected. The macrophages from collected cells were gated as CD64^+^ F4/80^+^, and further gated into transferred cells (CD64^+^ F4/80^+^ CD45.1^+^ CD45.2^-^); local AMs (CD64^+^ F4/80^+^ CD45.1^-^CD45.2^+^ Siglec F^+^), and local IMs (CD64^+^ F4/80^+^ CD45.1^-^CD45.2^+^ Siglec F^-^). (**B**). The WGA MFI of transferred cells, local AMs and local IMs collected from the digested lungs at day 7 (**C**) and day 14 (**D**). The WGA MFI of transferred cells and local AMs collected from the BAL fluid at day 7 (**E**) and day 14 (**F**). Representative flow plots of the transferred macrophages on the expression of Siglec-F on days 7 and 14 (**G**). Representative histograms of Siglec-F on days 7 and 14 (**H**). The transferred cells were divided into two subgroups based on the expression of Siglec-F. The MFI of WGA lectin staining from Siglec-F^+^ and Siglec-F^-^ transferred cells collected from the digested lung (**I**), and the BAL fluid (**J**). Each dot represents an individual mouse, data are presented as mean ± SEM and were analysed using unpaired t-tests. *, P ≤ 0.05; **, P ≤ 0.01; ***, P ≤ 0.001; ****, P ≤ 0.0001.

## Discussion

The differential expression of glycocalyx-associated components on specific macrophage subsets and macrophages in different anatomical locations is interesting. We used four different strategies to show that macrophage expression of the glycocalyx depends on where they reside in the lung (Figure 8). First, confocal imaging shows dense glycocalyx on macrophages in the airspaces that is absent on those in the interstitium. Second, bone marrow mesenchymal stem cells differentiated into macrophages with GM-CSF (highly expressed in the airspaces^30^) have higher RNA transcripts for glycocalyx synthesis proteins and their regulators. Further polarisation of GM-CSF monocyte-derived macrophages with IL-4 drives even greater expression of RNA and protein for glycocalyx components. Third, glycocalyx level increases on interstitial macrophages at the later stages of an influenza infection, at a time when virus is cleared and the repair process is beginning^31^. Fourth, glycocalyx-deficient macrophages instilled intranasally develop a glycocalyx in the airways over time.

**Figure 8:**
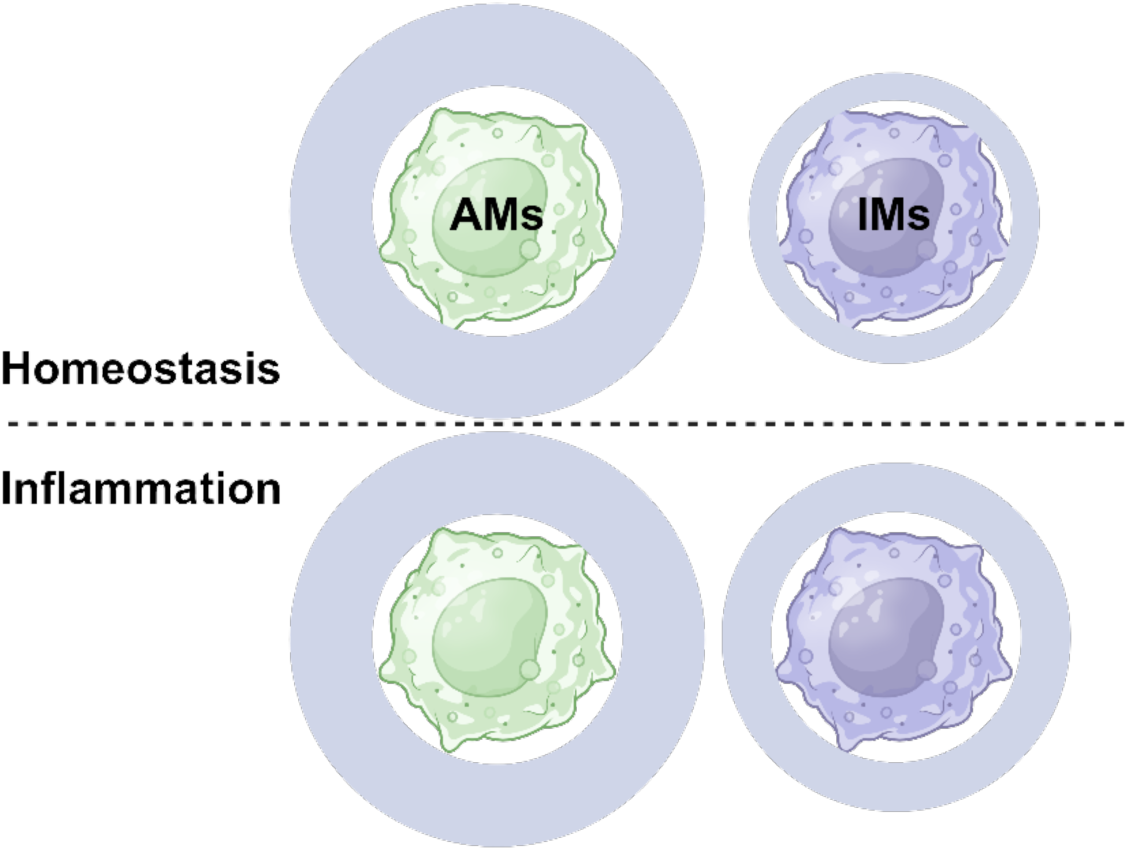
Airway macrophage specific expression of a dense glycocalyx and its remodelling following viral infection. Pulmonary macrophage glycocalyx is determined by anatomical location and inflammation. Airway macrophages are decorated with high levels of glycocalyx in both homeostasis and inflammation. Interstitial macrophages show low glycocalyx levels in homeostasis. During Influenza virus-induced inflammation, interstitial macrophages increased the glycocalyx levels.

There is indirect, but not direct, support for a macrophage glycocalyx in the literature. For example, HS and CS attach to approximately 40 core proteoglycans, including perlecan, agrin, Collagen XVIII and SDCs, amongst others^32^. SDC-1 is expressed on macrophages in rheumatoid arthritis^33,34^ and on human primary macrophages^35^, and is likely to be decorated with HS and CS. Human macrophages polarised to an M2/regulatory phenotype also express higher SDC-1^36^. Furthermore, cell-specific targeting of glycosaminoglycan chemical modification (sulfation) has been shown to affect macrophage function in other contexts^37,38^. Thus, evidence of macrophage SDC-1 expression provides the possibility for tethering of a glycocalyx, and the data corroborate the M2 bias of expression with the mRNA and protein we describe here.

The specific reduction of multiple glycocalyx components on M1 macrophages in our study may reflect reduced production or enhanced degradation. Exostatin 2 and 3 are involved in HS biosynthesis that occurs in the Golgi and the Golgi-endoplasmic reticulum interface^39,40^. Biosynthesis of HS is blocked by knockdown of *ext2* and ext2-like^41^. Furthermore, TNF and IL-1 activate sheddases that cleave the glycocalyx^42^, including from epithelial cells in acute respiratory distress syndrome^43^ and also immune cells themselves^11^, providing a source of biomarkers. Neuraminidase 2 also causes glycocalyx shedding^44^, and MMP9 liberates SDC-1 and −4 from human macrophages ^45^.

This raises the question of why it would be beneficial to reduce glycocalyx on macrophages, specifically in M1-dominated conditions. We hypothesise that sensing the glycocalyx through surface receptors triggers a functional programme of repair, and its absence may allow inflammation to proceed. This idea is supported by studies showing that attenuation of M1 macrophage polarisation and augmentation of M2 prevents glycocalyx shedding via a decrease in ADAM17 in a murine model of acute lung injury^46^. SDC-1 deletion, which, though not tested, would limit HS and CS binding, increases inflammation in murine asthma models^47,48^. Deficiency of EXTL2 (that drives HS production) promotes inflammation via TLR4^49^ and a variant of EXTL2 is associated with asthma exacerbation^50^. Again, the glycocalyx in these studies was not tested. Inflammation also accompanies a reduction of CS in mice^51^. Administration of CS reduces cytokines in fungal infection^52^, CpG-induced IL-6 in a macrophage cell line^53^ and peritoneal fibrosis via suppression of NF-κB^54^ suggesting that reduced glycocalyx promotes inflammation. Limitation of sheddases (that would cleave glycocalyx tethers) also attenuates the inflammatory environment in sepsis^55^. The predominant focus of these studies is the glycocalyx on epithelium and endothelium. Our combined data, including *in vitro* polarisation and analysis of influenza virus recovery and repair, suggest that the dominant anti-inflammatory activity of the glycocalyx may be via modulation of macrophage inflammatory tone.

We used soluble TLR agonists for *in vitro* mechanistic studies, as a glycocalyx is reported to be necessary for endocytosis of viruses and bacteria and so in vivo models may have confounding issues. For example, proteoglycans, which rely on B4GALT7, Ext1 and Ext2 for the attachment of HS and CS, are implicated in the endocytosis of influenza virus^56,57^. Furthermore, CRISPR-Cas9 deletion of GAG biosynthesis regulators, including B4GALT7, reduced the uptake of *Mycobacterium absessus* in macrophages, possibly by modulation of integrin accessibility^58^.

It is possible that modulation of inflammatory mediator production by macrophages following glycocalyx shedding may result from the unshielding of activating receptors^59^. However, the glycocalyx, when intact, may provide tonic signals to macrophages, limiting their inflammation, much in the same way that lung surfactant proteins reduce macrophage endocytosis and TLR signalling^22^. In general, receptor binding of individual glycocalyx components promotes an anti-inflammatory state in macrophages^48^. CS, for example, limits NF-κB^60–62^, reduces IL-6, NOS-2 and PGE2 synthase in bone marrow-derived macrophages^63,64^ and is reported to block LPS binding to CD44^65^. Furthermore, high molecular weight HA reduces inflammation upon binding to CD44^66^. However, these effects are equally likely to increase the macrophage glycocalyx, which was not tested in these prior studies. High heparanase activity is observed in COVID-19 and is associated with increased disease severity and pro-inflammatory IL-6, TNF, IL-1 and CCL2 production^67^. These effects are reversed by heparanase blockade^68^ though the impact on glycocalyx is untested In addition, deletion of SDC-1 (covalently linked to HS and CS^69^) enhances inflammation in an asthma model^47^. Whether these manipulations restore negative regulation and/or activating receptor shielding is unknown. However, our data provides evidence that macrophages may sense health, and when to inflame, through their glycocalyx.

In summary, macrophages express genes involved in glycocalyx biosynthesis, which suggests components are not simply adsorbed from the local environment. Differential regulation of glycocalyx density on macrophages in environmentally exposed sites makes sense, as does a degradation of the glycocalyx to signal a switch away from a healthy environment to inflammation. Regaining glycocalyx on macrophages is likely important to switch on repair programmes and also prevent the development of autoimmunity upon efferocytosis of apoptotic cells and the removal of host cell debris. Manipulation of glycocalyx components, therefore provides a strategy to increase or decrease the onset of inflammation.

## METHODS

### Animals

All mice (10-12 weeks old) were housed in the Biological Services Facility at 22 °C to 24 °C, with 45 % to 55 % humidity and a 12:12-h light-dark schedule. All procedures were approved by the University of Manchester Animal Welfare and Ethical Review Body and the Home Office UK in accordance with the UK Animals (Scientific Procedures) Act, 1986. B6.SJL-Ptprca Pepcb/BoyJ mice (CD45.1^+^ CD45.2^-^) were provided by Dr Grace Mallett at the University of Manchester. All C57BL/6J mice (CD45.1^-^CD45.2^+^) were purchased from ENVIGO UK Ltd.

#### Influenza infection

Mice were anaesthetised with isoflurane and intranasally infected with 30µL containing 5 plaque-forming units (PFU) of influenza A virus, Puerto Rico/8/34 (PR8), H1N1 or the same volume of PBS as a control. Mouse weight was collected daily and euthanised at day 7 or day 14 by intraperitoneal injection of 5 mg pentobarbitone. Bronchoalveolar lavage was conducted by instilling and recovering 5 ml of Hanks’ Balanced Salt solution (HBSS), with 0.5 mM EDTA (MERCK, 324506), from the lungs via an intratracheal cannula. The remaining lung tissue was resected. The left lobes (superior and inferior) were fixed in formalin and processed for paraffin wax embedding. The remaining right (superior, middle and inferior) lobes were homogenised and digested in 1 ml HBSS with Liberase TM (100 µg/ml; Millipore Sigma, 5401127001) and DNase I (50 ug/ml; Sigma, 11284932001) in HBSS, on a shaker, at 37 °C for 30 minutes. The digestion was stopped with 4 ml of HBSS containing 2% FBS and 2 mM EDTA (Thermo Fisher Scientific, AM9260G). The tissue was passed through a 70 µm sieve, washed with HBSS, and collected by centrifugation at 500 g for 5 minutes. Red blood cells were removed by incubating with 3ml of lysis buffer (Sigma-Aldrich, R7757) for three minutes at RT. After washing, cell viability was assessed using methyl orange and viability was enumerated on an automated cell counter (ChemomeTec, NucleoCounter NC-250) and the ChemoMetec Nucleo NC-250 software (ChemomeTec).

#### Studies using bone marrow-derived cells

Inthe cell transfer experiment, CD45.2 mice were given CD45.1 BMDMs to allow cell tracking. Briefly, the fibula and tibia bones of B6.SJL-Ptprca Pepcb/BoyJ mice (CD45.1^+^ CD45.2^-^) were flushed with to liberate bone marrow, red blood cells were lysed, and macrophages were derived from monocytes using 20 ng/ml M-CSF, in RPMI 1640 Medium (Thermo Fisher Scientific, 11879020) with 20% FBS (Gibco, 11533387), 100 U/ml penicillin (Gibco, 15140122), and 100 µg/ml streptomycin (Gibco, 15140122), and 20 mM HEPES solution (Sigma-Aldrich, H0887)for 7 days. The cell numbers were counted, and 1×10^6^ of the M-CSF-derived BMDMs was instilled intranasally into each C57BL/6J mice (CD45.1^-^CD45.2^+^) mouse with 25 µl PBS. The lungs were collected 7 and 14 days later.

For *in vitro* studies, bone marrow cells were cultured with 20 ng/ml recombinant murine M-CSF (PeproTech, 315-02) or 20 ng/ml recombinant murine GM-CSF (PeproTech, 315-03), in RPMI 1640 Medium mixture mentioned above. On day 4, media was refreshed with 20 ng/ml M-CSF or 20 ng/m GM-CSF and on day 8, non-adherent cells were removed. Adherent cells were recovered by scraping and centrifugation. In some experiments, BMDM were further polarised for 24 hours with the RPMI 1640 Medium mixture with 20 ng/ml recombinant murine IFN-γ (PeproTech, 315-05), 10 ng/ml Lipopolysaccharide (Invivogen, tlrl-eblps); or 20 ng/ml recombinant murine IL-4 (PeproTech, 214-14), before scraping. In all polarisations, cells were subjected to flow cytometric analysis. In some experiments where the RNA transcriptome was analysed, polarised adherent cells were incubated in 350 µl of RLT buffer (Qiagen, 74004) mixed with 1:100 beta-mercaptoethanol (Sigma-Aldrich, 60-24-2) to produce cell lysis and stored at −80°C.

### Flow cytometry staining and analysis

Approximately 2×10^6^ cells were incubated with mouse FcR Blocking Reagent (1:500, Miltenyl Biotec, Germany, 5210502523) and zombie UV (1:1000, BioLegend, USA, 423107) in 50µl PBS for 15 minutes at RT. After 50 µl of 2 % paraformaldehyde fixation at RT for 10 minutes, cells were stained with the indicated directly conjugated or unconjugated (followed by a secondary detection antibody) (**table S1)** antibodies in 50 µl PBS with 1% FBS, overnight at 4°C. After washing and fixation with 50 µl 2% paraformaldehyde samples were run on a BD FAC Fortessa Flow Cytometer (BD Bioscience, UK), using the FACS Diva software (BD Bioscience, Belgium) and analysed with the FlowJo software (Tree Star, USA).

### Confocal microscopy

The formalin fixed and paraffin embedded lung tissues were trimmed and attached on glass slides (MERCK, CLS294875X25). The slides were deparaffinized with xylene and ethanol. Antigen retrieval was accomplished at 95 °C for 20 minutes in 400 ml Tris-EDTA buffer (Sigma-Aldrich, T9285). Slides were then permeabilised with 200 ml 0.5 % Triton X-100 (Sigma-Aldrich, X100) and blocking buffer applied (1% FBS, 2% donkey serum (MERCK, D9663) and 0.05% Tween-20 (MERCK. P2287)) to cover the tissue on the glass slides for 30 minutes. Primary antibodies were applied to cover the tissue overnight at 4°C, and if required, followed by secondary antibodies at RT for 1 hour (**tables S2)**. The slides were stained with equivalent amount of 0.2 μg/ml 4ʹ,6-diamidino-2-phenylindole (DAPI, Thermo Scientific, 62248) in H^2^O for 5 minutes. The slides were washed with 200 ml 0.1% Tween 20 (Sigma-Aldrich, 11332465001) in PBS and two washes with 200 ml PBS after each step. The slides were mounted with ProLong™ Gold Antifade Mountant (Invitrogen, P36934), and covered with cover slides (Corning, CLS2975224). Slides were imaged using an EVOS FL Auto imaging system (Thermo Fisher Scientific) and analysed using Image-J Software.

### RNA isolation, reverse transcription, and RT-qPCR

RNA was isolated using the RNeasy Micro Kit (Qiagen, 74004) according to the manufacturer’s instructions. The total RNA concentration was quantified with a NanoDrop 2000/2000c Spectrophotometer (Thermo Fisher Scientific, ND-2000). Reverse transcription was conducted using the High-Capacity RNA-to-cDNA Kit (Applied Biosystems, 4387406) and 100 ng RNA was used in each reverse transcription. The samples were incubated at 37 °C for 60 minutes, then at 95 °C for 5 minutes for heat inactivation and stored at 4 °C. After reverse transcription, the cDNA was diluted 1:4 in nuclease-free water. Reactions were performed in triplicate on a MicroAmp™ Optical 384-Well Reaction Plate with Barcode (Applied Biosystems, 4309849). Each qPCR reaction contained 5 μl PowerUp SYBR Green Master Mix (Applied Biosystems, A25742), 0.6 μl 10 μM each of forward primer and reverse primer, 2.4 μl ddH2O and 2 μl cDNA to perform a 10-μl reaction volume. was analysed on a 7900HT Fast Real-Time PCR System (Applied Biosystems). The PCR cycle included 95 °C for 5 minutes, 40 x 15 seconds at 95 °C, 20 seconds at 57 °C and 20 seconds at 72 °C, followed by 95 °C, 60°C and 95°C for 15 seconds each, sequentially. Relative mRNA levels were calculated compared to the average mRNA expression of the housekeeping gene beta-2-microglobulin (B2m). Relative mRNA expression and fold changes were calculated based on the ΔΔCT method. Ct, ΔCt, ΔΔCt and 2-ΔΔCt were calculated (Livak & Schmittgen, 2001). Primers: KiCqStart™ Primers purchased from MERCK: *b2m* (M_B2m_1), *nos-2* (M_Nos2_1), *il-6* (M_Il6_1), *thf-α* (M_Tnf_1), *chi3l-1* (M_Chi3l1_1), *b4galt-7* (M_B4galt7_1), *ext-2* (M_Ext2_1), *extl-1* (M_Extl1_1), *extl-2* (M_Extl2_1), *extl-3* (M_Extl3_1), *sdc-1* (M_Sdc1_1), *sdc-2* (M_Sdc2_1), *sdc-3* (M_Sdc3_3), *sdc-4* (M_Sdc4_1), *has-1* (M_Has1_1), *has-2* (M_Has2_1), *has-3* (M_Has3_1), *mmp-9* (M_Mmp9_1), *mmp-12* (M_Mmp12_1), and *mmp-13* (M_Mmp13_1).

### Statistical analysis

All statistics were calculated using GraphPad Prism 10.2.0 (GraphPad Software Inc., La Jolla, CA). When comparing between two groups, an unpaired *t-test* was performed to determine statistical significance. When comparing among three or more groups, an analysis of variance (ANOVA) was performed. Error bars indicate SEM.

## Acknowledgements

TH and ZZ are supported by The Wellcome Trust (202865/Z/16/Z). Sir Henry Dale fellowship jointly funded by the Wellcome Trust and Royal Society 218570/Z/19/Z (DPD). Wellcome Trust center grant 203128/A/16/Z (DPD).

