## Supplementary information for "Airway macrophage specific expression of a dense glycocalyx and its remodelling following viral infection"

**Table S1: Reagents and antibodies for flow cytometry staining.** *MERCK, ASMBIO, # MBL International

| Reagent | Clone | Host/Isotype | Dilution | Conjugate | Catalogue |
|---|---|---|---|---|---|
| *Wheat germ agglutinin (WGA) | | | 1:2000 | FITC | L4895 |
| Anti-mouse CD45 | 30-F11 | Rat IgG2b, κ | 1:200 | Brilliant Violet 510 | 103137 |
| Anti-mouse Ly-6G | 1A8 | Rat IgG2a, κ | 1:200 | APC/Cyanine7 | 127624 |
| Anti-mouse CD19 | 6D5 | Rat IgG2a, κ | 1:200 | APC | 115529 |
| Anti-mouse/ human CD11b | M1/70 | Rat IgG2b, κ | 1:500 | Brilliant Violet 711 | 101242 |
| Anti-mouse CD11c | N418 | Armenian Hamster IgG | 1:400 | Brilliant Violet 605 | 117334 |
| Anti-mouse CD64 (FcγRI) | X54-5/7.1 | Mouse IgG1, κ | 1:200 | Brilliant Violet 421 | 139309 |
| +Heparan Sulfate | F58-10E4 | Mouse IgM, κ | 1:200 | Biotin | 10E4260917_060120 |
| Anti-mouse NK-1.1 | PK136 | Mouse IgG2a κ | 1:200 | BV605 | 108753 |
| Anti-mouse Ly-6C | HK1.4 | Rat IgG2c, κ | 1:200 | Alexa Fluor 700 | 128023 |
| Anti-mouse CD138 | 281-2 | Rat IgG2a, κ | 1:200 | PE/Cyanine 7 | 142513 |
| Anti-mouse CX3CR1 | QA16A03 | Mouse IgG1, κ | 1:200 | APC | 153707 |
| Anti-mouse CD3 | 17A2 | Rat IgG2b, κ | 1:200 | FITC | 100203 |
| Anti-mouse CD170 (Siglec-F) | S17007L | Rat IgG2a, κ | 1:200 | PE | 155505 |
| Anti-mouse CD4 | GK1.5 | Rat IgG2b, κ | 1:200 | PE/Cyanine 5 | 100409 |
| #H-2Db Influenza NP Tetramer<br>ASNENMETM | | | 1:200 | BV421 | TB-M508-4 |
| Anti-mouse CD8a Antibody | 53-6.7 | Rat IgG2a, κ | 1:200 | BV650 | 100742 |

| | | | | | |
|---|---|---|---|---|---|
| Anti-mouse CD64 (FcγRI) | X54-5/7.1 | Mouse IgG1, κ | 1:200 | PE/Cyanine 7 | 139313 |
| Anti-mouse CD284 (TLR4) | SA15-21 | Rat IgG2a, κ | 1:200 | PE | 145403 |
| Anti-mouse /human CD44 | IM7 | Rat IgG2b, κ | 1:200 | PE/Cyanine 5 | 103009 |
| Anti-mouse CD45.1 | A20 | Mouse (A.SW) IgG2a, κ | 1:200 | Brilliant Violet 650 | 110736 |
| Anti-mouse CD45.2 | 104 | Mouse (SJL) IgG2a, κ | 1:200 | Brilliant Violet 785 | 109839 |
| Streptavidin | | | 1:200 | Alexa Fluor 594 | 405240 |
| Streptavidin | | | 1:200 | Alexa Fluor 647 | 405237 |

**Table S2: Reagents and dilutions for immunofluorescence staining**

| Reagent | Clone | Host/Isotype | Dilution | Conjugate | Catalogue |
|---|---|---|---|---|---|
| Wheat germ agglutinin (WGA) | | | 1:2000 | FITC | MERCK L4895 |
| CD64 Rabbit | 008 | Rabbit / IgG | 1:200 | Unconjugated | Invitrogen MA5-29705 |
| Rat Anti-Mouse Siglec-F | E50-2440 | Rat (LOU) IgG2a, κ | 1:200 | Unconjugated | 552125 BD Pharmingen |
| Rabbit IgG NorthernLights™ NL637 | | Polyclonal Donkey IgG | 1:200 | NorthernLights 637 | R&D System NL005 |
| Donkey anti-Rat IgG (H+L) Highly | | Polyclonal Donkey IgG | 1:200 | Alexa Fluor 594 | Invitrogen A21209 |

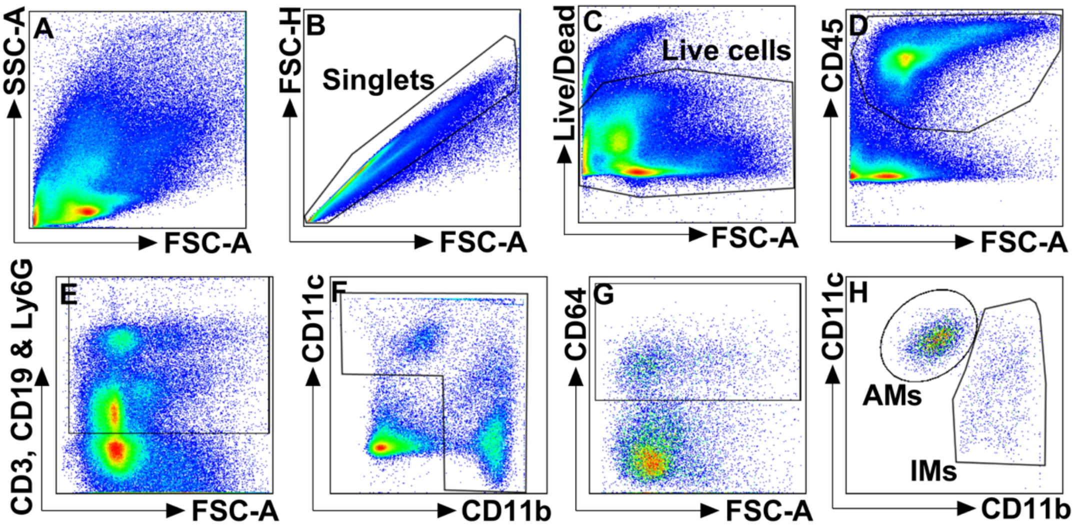


**Supplementary Figure 1: Gating strategy for lung myeloid cells.** Cells were isolated from the digested lungs of female C57BL/6 mice. After staining, the cells were run through a BD Fortessa flow cytometer (**A**) and singlets were identified (**B**). Live cells (**C**) and $CD45^{+}$ cells (**D**) were gated. Cells positive for Ly6G, CD3, or CD19 were gated out to remove T cells and B cells (**E**). Cells either CD11c or CD11b positive were gated (**F**), followed by $CD64^{+}$ cells to identify macrophages (**G**). The remaining cells with $CD11b^{high}$ were classified as IMs, and those with $CD11c^{high}$ were classified as AMs (**H**).

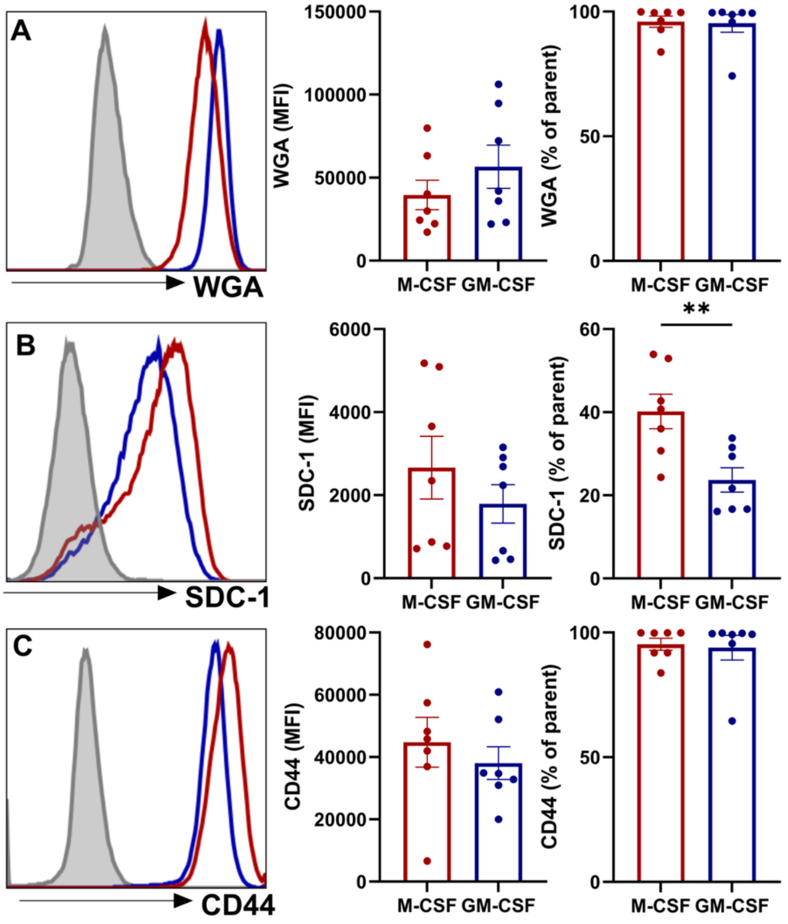


**Supplementary Figure 2: Flow cytometry comparison of the glycocalyx on unpolarized M-CSF-derived BMDMs and GM-CSF-derived BMDMs.** The femurs of C57BL/6 female mice were flushed with HBSS to collect bone marrow cells. The cells were then cultured in 20 ng/ml M-CSF or GM-CSF to derive the unpolarized (M0) M-CSF-derived BMDMs and GM-CSF-derived BMDMs. The medium fluorescence intensity (MFI) and percentage of cells positive for WGA lectin (detects the N-acetyl-glucosamine) (**A**), SDC-1 (**B**), and CD44 (**C**). Data are collected from two independent experiments with three and four repeats, presented as the mean ± SEM. Representative histograms are shown: M-CSF-derived BMDMs (red), GM-CSF-derived BMDMs (blue), and FMO control (grey). Data was analysed with unpaired t-tests. *, $P \leq 0.05$; **,$P \leq 0.01$.

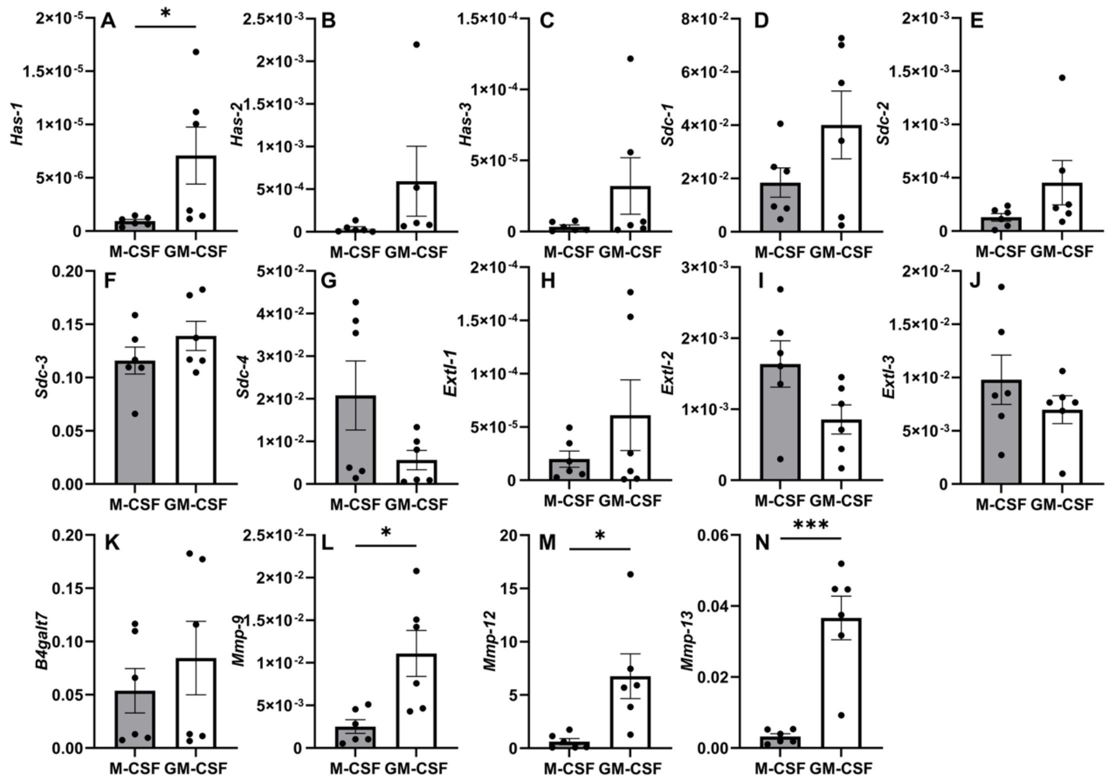


**Supplementary Figure 3: GM-CSF-derived BMDMs express higher levels of genes for glycocalyx degradation than M-CSF-derived BMDMs.** Bone marrow cells were derived with M-CSF or GM-CSF. The glycocalyx-relative mRNA expression was measured in GM-CSF- and M-CSF-differentiated BMDMs. The qPCR data are normalised to the housekeeping gene *b2m* and displayed as the relative expression levels in comparison to the expression level of *b2m*. *Has-1* (**A**), *has-2* (**B**), *has-3* (**C**), *sdc-1* (**D**), *sdc-2* (**E**), *sdc-3* (**F**), *sdc-4* (**G**), *extl-1* (**H**), *extl-2* (**I**), *extl-3* (**J**), *b4galt7* (**K**), *mmp-9* (**L**), *mmp-12* (**M**), and *mmp-13* (**N**) Data are representative of two independent experiments, each with three technical repeats, shown as the mean ± SEM. Data was analysed with unpaired t-tests. *, $P \leq 0.05$; **, $P \leq 0.01$; ***, $P \leq 0.001$.

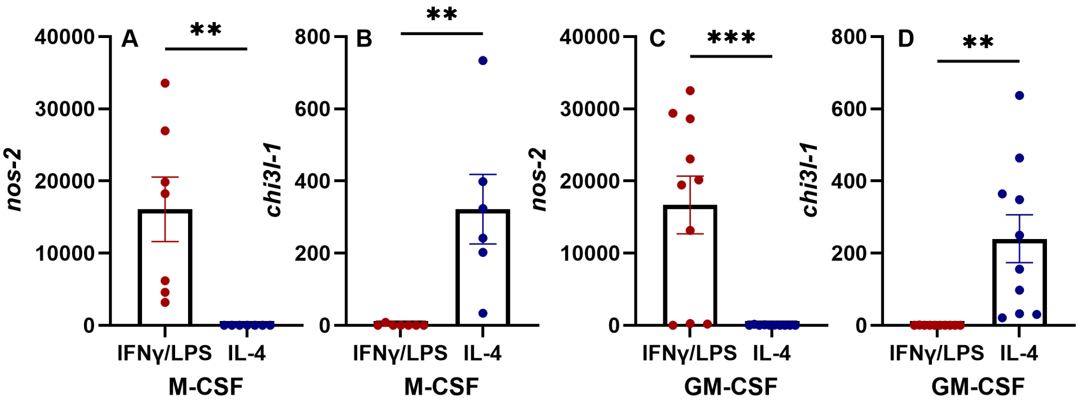


**Supplementary Figure 4**: **Expression of M1 and M2 polarisation markers in BMDMs.** The mRNA expression of *nos-2* (**A**) and *chi3l-1* (**B**), display as fold change over the level of unpolarised M0, in M-CSF-derived BMDMs. Data are representative of two independent experiments. The mRNA expression of *nos-2* (**C**) and *chi3l-1* (**D**), display as fold change over the level of unpolarised M0, in GM-CSF-derived BMDMs. Data are representative of three independent experiments. Each experiment has three technical repeats, shown as the mean ± SEM. Data was analysed with unpaired t-tests. **, $P \leq 0.01$; ***, $P \leq 0.001$.

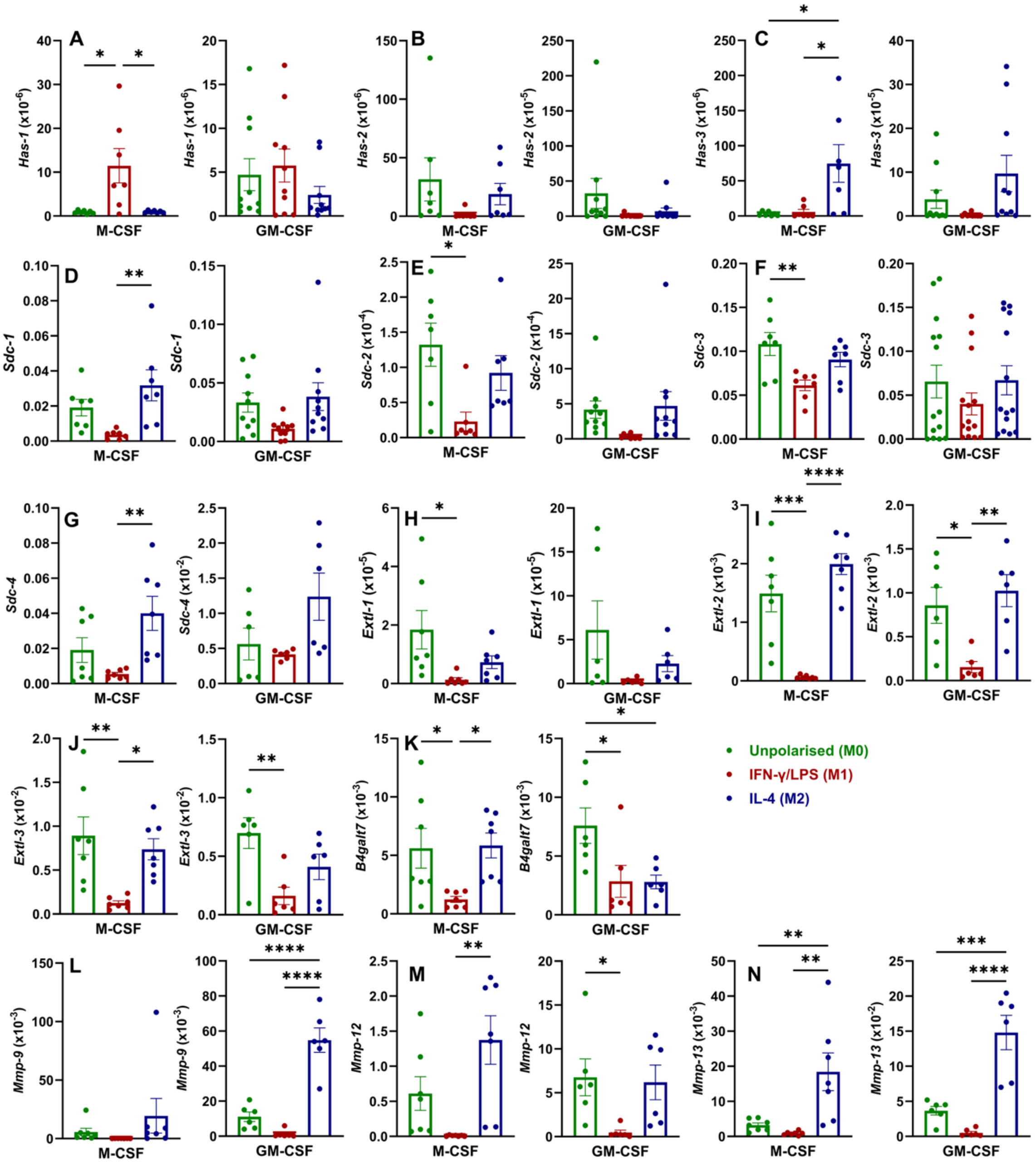


**Supplementary Figure 5**: **Relative expression levels of glycocalyx-related genes in M-CSF- and GM-CSF-derived BMDMs in M1 and M2 polarising stimuli.** M-CSF-derived BMDMs and GM-CSF-derived BMDMs were left unstimulated (green) or polarized to classically (M1) (red) and alternatively activated (M2) (blue) macrophages. The relative mRNA expression levels were measured according to the housekeeping gene *b2m*. *Has-1* (**A**), *has-2* (**B**), *has-3* (**C**), *sdc-1* (**D**), *sdc-2* (**E**), *sdc-3* (**F**), *sdc-4* (**G**), *extl-1* (**H**), *extl-2* (**I**), *extl-3* (**J**), *b4galt7* (**K**), *mmp-9* (**L**), *mmp-12* (**M**), and *mmp-13* (**N**). Data are representative of two or three independent experiments with three technical repeats, as the mean ± SEM. Data was analysed with unpaired t-tests. *, P ≤ 0.05; **, P

≤ 0.01; ***, P ≤ 0.001; ****, P ≤ 0.0001.

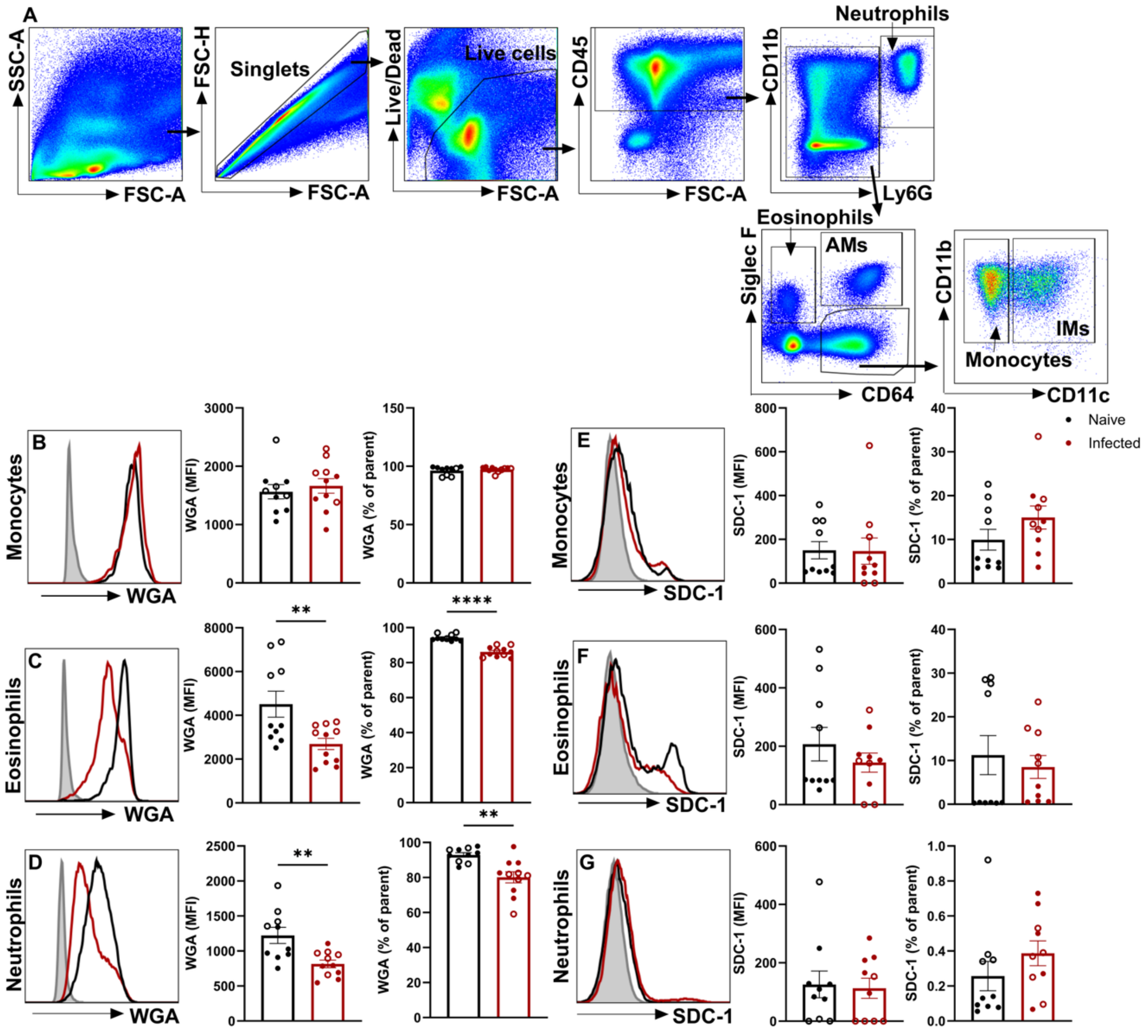


**Supplementary Figure 6: SDC-1 levels remain unchanged following the flu infection on monocytes, eosinophils and neutrophils**. At day 0, mice were intranasally given 5 PFU IAV PR8 in 25 ul PBS, or given the same volume of PBS (25 µl) as control. On day 7, the lung tissue from both groups was harvested. Myeloid cells were distinguished by a flow cytometer, including neutrophils ($CD45^+$ $Ly6G^{high}$ $CD11b^+$), airway macrophages ($CD45^+$ $Ly6G^{medium/low}$ $CD64^+$ Siglec $F^+$), eosinophils ($CD45^+$ $Ly6G^{medium/low}$ $CD64^-$ Siglec $F^+$), IMs ($CD45^+$ $Ly6G^{medium/low}$ $CD64^+$ Siglec $F^-$ $CD11b^+$ $CD11c^+$), and monocytes ($CD45^+$ $Ly6G^{medium/low}$ $CD64^+$ Siglec $F^-$ $CD11b^+$ $CD11c^-$) (**A**).The MFI, and percentage, of general glycocalyx showed by WGA lectin staining of monocytes (**B**), eosinophils (**C**), and neutrophils (**D**). The MFI and percentage of monocytes (**E**), eosinophils (**F**), and neutrophils (**G**) positive for SDC-1 expression. The histograms are representative of one mouse per group. Data are presented as mean ± SEM and was analysed by unpaired t-test. *, $P \leq 0.05$; **, $P \leq 0.01$; ***, $P \leq 0.001$; ****, $P \leq 0.0001$. Data are representative of two independent experiments, with n = 10 or 11 in each experiment. Data from the two experiments were distinguished by the shape of the points.
